# Expression and pharmacological inhibition of TrkB and EGFR in glioblastoma

**DOI:** 10.1101/2020.02.03.932608

**Authors:** Kelly de Vargas Pinheiro, Amanda Thomaz, Bárbara Kunzler Souza, Victoria Anne Metcalfe, Natália Hogetop Freire, André Tesainer Brunetto, Caroline Brunetto de Farias, Mariane Jaeger, Victorio Bambini, Christopher G.S. Smith, Lisa Shaw, Rafael Roesler

**Affiliations:** Cancer and Neurobiology Laboratory, Experimental Research Center, Clinical Hospital (CPE-HCPA), Federal University of Rio Grande do Sul, 90035-003 Porto Alegre, RS, Brazil; Department of Pharmacology, Institute for Basic Health Sciences, Federal University of Rio Grande do Sul, 90050-170 Porto Alegre, RS, Brazil; Children’s Cancer Institute, 90620-110 Porto Alegre, RS, Brazil; School of Pharmacy and Biomedical Sciences, Faculty of Clinical and Biomedical Sciences, University of Central Lancashire, Preston, Lancashire, PR1 2HE, United Kingdom

**Author notes:** Correspondence: Rafael Roesler, Department of Pharmacology, Institute for Basic Health Sciences, Federal University of Rio Grande do Sul, Rua Sarmento Leite, 500 (ICBS, Campus Centro/UFRGS), 90050-170 Porto Alegre, RS, Brazil. Division of Biomedical and Life Sciences, Faculty of Health and Medicine, Lancaster University, Lancaster LA 4YG, United Kingdom.

**Keywords:** Brain tumor, Epidermal growth factor receptor, Glioblastoma, Growth factor receptor, Neurotrophin, Tropomyosin receptor kinase B

## Abstract

**Background:** A member of the Trk family of neurotrophin receptors, tropomyosin receptor kinase B (TrkB, encoded by the *NTRK2* gene) is an increasingly important target in various cancer types, including glioblastoma (GBM). *EGFR* is among the most frequently altered oncogenes in GBM, and EGFR inhibition has been tested as an experimental therapy. Functional interactions between EGFR and TrkB have been demonstrated. In the present study, we investigated the role of TrkB and EGFR, and their interactions, in GBM.

**Methods and Results:** Analyses of *NTRK2* and *EGFR* gene expression from The Cancer Genome Atlas (TCGA) datasets showed an increase in *NTRK2* expression in the proneural subtype of GBM, and a strong correlation between *NTRK2* and *EGFR* expression in glioma CpG island methylator phenotype (G-CIMP+) samples. We showed that when TrkB and EGFR inhibitors were combined, the inhibitory effect on A172 human GBM cells was more pronounced than when either inhibitor was given alone. When U87MG GBM cells were xenografted into the flank of nude mice, tumor growth was delayed by treatment with TrkB and EGFR inhibitors, given alone or combined, only at specific time points. Intracranial GBM growth in mice was not significantly affected by drug treatments.

**Conclusions:** Our findings indicate that correlations between *NTRK2* and *EGFR* expression occur in specific GBM subgroups. Also, our results using cultured cells suggest for the first time the potential of combining TrkB and EGFR inhibition for the treatment of GBM.

## Introduction

Growth factor receptors constitute many current and potential targets for molecularly specific therapies in cancer. The epidermal growth factor receptor (EGFR), a member of the ERBB family of transmembrane receptor tyrosine kinase (RTK) family, frequently shows gene amplification and activating mutations that contribute to driving the growth of lung and colorectal cancers [1]. EGFR is the target of clinically used small molecule inhibitors including erlotinib and gefitinib and monoclonal antibodies including cetuximab and panitumumab [2].

*EGFR* is among the most frequently altered oncogenes in glioblastoma (GBM), with 57% of tumors analyzed by the The Cancer Genome Atlas Research Network (TCGA) showing amplification, mutation, rearrangement, or altered splicing [3]. GBM is the most aggressive type of primary malignant brain tumor. Current treatment based on combining surgical resection followed by radiotherapy and chemotherapy results in a median overall survival of less than 2 years [4, 5]. To date, clinical trials with EGFR inhibitors in patients with GBM have failed to successfully improve outcomes [6, 7]. A proposed experimental strategy to reduce resistance and improve effectiveness has been to combine EGFR inhibitors with other targeted agents acting on pathways that crosstalk with EGFR signaling [8, 9].

EGFR has been shown to functionally interact with tropomyosin receptor kinase B (TrkB, encoded by the *NTRK2* gene), a member of the Trk family of neurotrophin receptors. Neurotrophins are secreted proteins importantly involved in central nervous system development, neurogenesis, and neuronal survival and plasticity [10]. Increasing evidence indicates that cancers can hijack neurotrophin signaling systems to promote tumor progression and resistance to treatment [9, 11]. In colorectal cancer cells, TrkB activation by its endogenous ligand, brain-derived neurotrophic factor (BDNF), promotes resistance against the EGFR inhibitor cetuximab, whereas co-treatment with TrkB and EGFR inhibitors reduce cell viability [12].

GBM tumors express BDNF and TrkB, and TrkB activation enhances the viability of brain tumor stem cells (BTSCs) from human GBMs, whereas its inhibition reduces BTSC growth [13]. TrkB blockade also hinders viability of cultured human GBM cells [14]. BDNF secreted by more differentiated GBM cells supports the growth of TrkB-expressing GBM BTSCs [15]. *NTRK* gene fusions are currently established as oncogenic drivers of various adult and pediatric tumor types, and larotrectinib, a first-in-class small molecule Trk inhibitor, has received approval for patients with solid tumors harboring *NTRK* fusions [16, 17]. A gene fusion involving *NTRK2* has been found to confer distinctive morphology and an aggressive phenotype in a case of low-grade glioma [18].

Here, we investigated the role of TrkB and EGFR in GBM. First, we show data on the expression of *NTRK2* and correlations with *EGFR* expression in GBM tumors from The Cancer Genome Atlas (TGCA) datasets. We then went on to examine the effects of inhibiting EGFR and TrkB receptors, either alone or in combination, in experimental models of GBM.

## Materials and methodsxs

### *NTRK2* and *EGFR* expression profiling in TCGA GBM datasets

*NTRK2* expression levels were examined in previously described The Cancer Genome Atlas (TCGA *n=* 631 samples) GBM dataset and TCGA Lower Grade Glioma and Glioblastoma (n= 702 samples) dataset. We used the data from gene expression array (platform: AffyU133a, version: 2017-09-08), gene expression RNAseq (platform: IlluminaHiSeq, version: 2017-09-08) and copy number (type: gene-level GISTIC2 threshold, version: 2017-09-08), data were obtained from the University of California–Santa Cruz Xena Public Data Hubs website at http://xena.ucsc.edu/. We analyzed the expression levels of NTRK2 regarding primary disease, primary tumor and recurrent samples, cytosine-phosphate-guanine (CpG) island methylator phenotype (G-CIMP), TCGA molecular subtypes, and the correlation with *EGFR* expression and amplification. Additionality, we evaluated the correlation between gene expression levels of *NTRK2* with patient survival outcomes.

### Cell culture

Human GBM cells A172 and U87MG were obtained from American Type Culture Collection (Rockville, MD, USA) and cultured in Dulbecco’s Modified Eagle’s Medium (DMEM) low glucose supplemented with 10% fetal bovine serum (FBS, Gibco® by Thermo Fisher Scientific, Life Technologies, Brazil), 1% penicillin/streptomycin and 0.1% fungizone® (250 mg/kg; Invitrogen Life Technologies, São Paulo, Brazil). Cells were maintained in a humidified atmosphere at 37°C and 5% CO_2_.

### Drug treatments

Selective antagonists of TrkB (ANA-12) and EGFR (Tyrphostin AG 1478) were obtained from Sigma Aldrich (St. Louis, MO). ANA-12 was diluted in dimethyl sulfoxide (DMSO) in a stock solution of 6140 µM and AG 1478 was diluted in ethanol (EtOH) in a stock solution of 3167 µM. Other chemical reagents were obtained from qualified national and international suppliers.

### Cytotoxicity

After confluence, A172 and U87MG cells were trypsinized, placed in 96-well plates at an initial density of 5.0 × 10^3^ cells per well and after 24h the medium was replaced by increasing concentrations of ANA-12 (0, 1, 10, 20, 30 and 50 µM), AG-1478 (0, 1, 5, 10, 20 and 30 µM) and also combinations of both inhibitors for 24, 48 or 72 h, while the control cells were maintained in DMSO or EtOH when the treatments were used alone or a vehicle solution (DMSO plus EtOH) when the treatments were used in combination. In any of the situations the vehicles used did not exceed the concentration of 1% (v/v). The effect on cell cytotoxicity was evaluated using the trypan blue exclusion method in the Neubauer chamber [12, 19]. All assays were performed in triplicate and repeated in three independent sets.

### Cell cycle

To assess cell cycle, GBM cells were cultured in 12-well plates under the same conditions as described above and after 24 h of exposure to treatments cells were detached, centrifuged and washed with PBS twice. The cells were then resuspended in 50 μg/ml propidium iodide (Sigma-Aldrich, St. Louis, Mo., USA) in 0.1% Triton X-100 in 0.1% sodium citrate solution and incubated on ice and protected from light for 15 min. The cells were analyzed by flow cytometry (Attune® Applied Biosystems) and 20,000 events were collected per sample. Three individual experiments were performed.

### Ethics statement

Experimental procedures for the subcutaneous xenograft GBM model were performed in accordance with the Brazilian Guidelines for the Care and Use of Animals in Research and Teaching [DBCA, published by National Council for the Control of Animal Experimentation (CONCEA), and approved by the institutional Animal Care Committee (Comissão de Ética no Uso de Animais CEUA, Hospital de Clínicas de Porto Alegre-HCPA), under protocol number 20160098. Animal experiments for the orthotopic xenograft GBM model were carried out under Animals (Scientific Procedures) Act, 1986 and in accordance to institutional guidelines.

### Animals and tumor xenografts

Balb/c nude mice (6 to 12 weeks old) were obtained from the University Hospital Animal Research Facility (UEA, CPE-HCPA) or from Charles River Laboratories. Animals were housed four per cage and kept under aseptic conditions in ventilated cages, maintained on a 12 h light/dark cycle at a room temperature of 22 ± 2°C. They were allowed *ad libitum* access to standardized pellet food and water. For the *ex-vivo* pharmacological inhibition, U87MG cells were cultured in 75 cm^2^ or 175 cm^2^ culture flasks and treated with 13.85 µM of ANA-12, 13.26 µM of AG 1478 (alone or in combination) or vehicle (DMSO plus EtOH) for 24 h. A total of 1 × 10^6^ viable cells were processed in serum-free DMEM and diluted 1:1 with Matrigel (Corning, Corning, USA) and then injected subcutaneously (s.c) into the right flank of nude mice (6-8 mice per group). Measurements started five days after cells inoculation, when tumors reached approximately 40-75 mm^3^. The dimensions, length (L) and width (W), of the resulting tumors were determined every two days using a manual caliper, and the tumor volume (mm^3^) was calculated using the formula: tumor volume = [length^2^ x width/2]. When tumors reached the endpoint (800-1000 mm^3^) mice were euthanized, and the tumors were excised, measured, and weighed.

For the orthotopic xenograft model, mice were anesthetized with isoflurane (4% oxygen) and placed in a stereotactic platform. The tops of the heads were disinfected, and a small incision was made in the scalp over the midline. A burr hole was made in the skull to a position 2 mm posterior and 1.5 mm lateral to the bregma in the right cerebral hemisphere. Next, mice were injected with 40,000 U87MG cells processed in serum-free DMEM in a volume of 2 µl using an injector syringe pump. The burr hole in the skull was closed with sterile bone wax and veterinary tissue glue was used to seal the incision. After surgery, the mice were placed in a recovery cage set to 37 °C until the animal recovered consciousness. Mice were monitored daily for signs of sickness, pain or weight loss. Seven days followed the surgery the animals were randomized into 4 groups (n=5 per group) to receive intraperitoneal (i.p.) injections for 21 days, and were treated by a blinded investigator with ANA-12 (1 mg/kg/daily plus vehicle for AG1478 every 3 days), AG1478 (10 mg/kg every three days plus vehicle for ANA-12 daily), ANA-12 (1 mg/kg daily) plus AG 1478 (10 mg/kg every three days). Control animals received ANA-12 and vehicle (DMSO 2% in saline solution 0.9%) daily and vehicle for AG1478 (DMSO 30µl every three days). Drug doses for *in vivo* experiments were chosen on the basis of previous studies [20 – 24]. Both drugs readily enter the brain when given systemically in mice or rats [21, 25].

### Fluorescence imaging

Administration of 5-aminolevulinic acid (5-ALA), that leads to the synthesis and accumulation of fluorescent protoporphyrin IX (PPIX), has been used with fluorescence-guided surgery to directly visualize high-grade gliomas. After 21 days of treatment, the animals received an i.p. injection of 5-ALA; 50 mg/kg and after 1 hour the mice were euthanized by cervical dislocation and the brains were removed to be analyzed. The IVIS system was used to record the ex-vivo fluorescent signal from tumors. First, we imaged the whole brains and next we used a brain matrix to cut sequential 1 mm slices, in order to improve fluorescence detection, and the slices were also imaged. Data acquisition and analysis was performed with Living Image software (Caliper LS living image version: 4.5.2.18424-september 11 2015, camera: IS1621N6980, Andor, iKon). In radiant efficiency mode, the wavelengths for emission and excitation were 620 and 420 respectively, and fixed size of regions of interest (ROIs) were drawn covering the whole tumor. The fluorescent signals (total radiant efficiency/sec) were quantitated by subtracting background fluorescence from tumor signal.

### Statistical analysis

Statistical analysis of gene expression in two or more groups from the TCGA transcriptome datasets were performed using Welch’s t-test and Kruskal-Wallis test for significance and Dunn’s tests for post hoc comparisons, respectively. Correlation analysis were performed by the Pearson correlation method. Survival distribution was estimated according to the Kaplan–Meier method using a median cut-off and log-rank statistics. These analyses were executed using the Graphpad 8.0 software. Significant differences were revealed by *p* values below 0.05.

Experimental data were expressed as mean ± SEM. *In vitro* and *in vivo* experiments were analyzed using the GraphPad Prism software package, v. 5.0 (GraphPad, San Diego, CA). The level of significance between different experimental groups was performed using analysis of variance (ANOVA) followed by appropriate post-hoc tests; *p* < 0.05 was considered statistically significant. A drug combination analysis was performed based on the 50% inhibitory concentration (IC50) of the treatments. For calculation of IC50 data were fitted in a dose response curve (Graphpad Prism v. 5.0) using the equation Y=100/(1+10^((X-LogIC50))). The interaction between ANA-12 and AG-1478 was assessed by the combination index method (Cl) [26]. Synergism, addition and antagonism between drug combinations were defined as CI < 0.9, CI = 0.9-1.1 and CI > 1.1, respectively.

## Results

### *NTRK2* expression levels are increased in lower grade glioma, proneural GBM subtype and GBM methylated phenotype in GBM patients

We evaluated TCGA GBM datasets to explore whether *NTRK2* expression correlates with *EGFR* expression, GBM subtype and patient survival. First, we classified TCGA samples by primary disease type between lower grade glioma (LGG) and GBM, based on array data available for 702 samples. *NTRK2* expression was increased in LGG in comparison to GBM samples (Fig. 1A, *p <* 0.0001). We also evaluated the levels of *NTRK2* across normal brain samples, primary and recurrent tumors, however, no statistical differences were observed between these groups (Fig. 1B, *p =* 0.220). Based on the definition of GBM molecular subtypes, we found that *NTRK2* expression was lower in the mesenchymal subtype when compared with neural and proneural subtypes (Fig. 1C, *p <* 0.01). *NTRK2* expression was highest in the proneural group in comparison with classical and mesenchymal subtypes (Fig. 1C, *p <* 0.0001). Given that the proneural group is associated with *IDH* mutations and *IDH*-mutant gliomas manifest the cytosine-phosphate-guanine (CpG) island methylator phenotype (G-CIMP), we evaluated the expression of *NTRK2* regarding the methylation phenotype between G-CIMP^+^ and G-CIMP-. We observed an increased expression of *NTRK2* in G-CIMP^+^ samples (Fig. 1D, *p <* 0.01), which is consistent with the increased expression in the proneural subtype.

**Fig 1.**
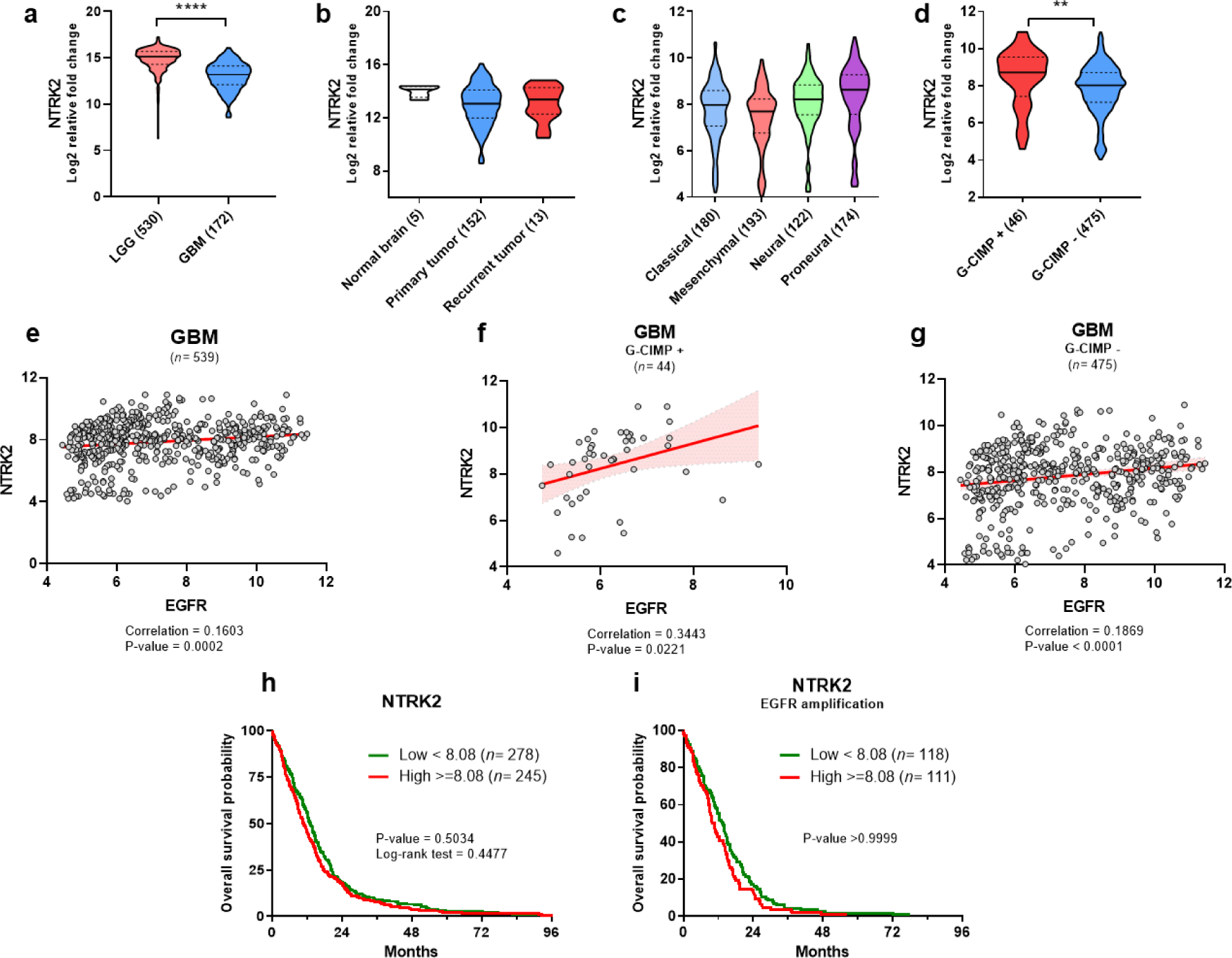
*NTRK2* expression and its correlation with *EGFR* expression levels in TCGA GBM datasets. Transcript levels of *NTRK2* were examined in previously described transcriptome datasets from the TCGA GBM dataset (*n* = 631 samples) and the TCGA Lower Grade Glioma (LGG) and GBM (*n* =702 samples) dataset. (A) Expression levels of *NTRK2* in LGG and GBM patient cohorts. (B) Normal brain, primary and recurrent GBM samples. (C) GBM subtypes. (D) cytosine-phosphate-guanine (CpG) island methylator phenotype (G-CIMP). Data was presented in violin plot format as log2-transformed signal intensity and statistical analyses were performed using Kruskal-Wallis test followed by Dunn’s post hoc tests, * *p* < 0.05; ** *p* < 0.01; *** *p* < 0.001 and **** *p* < 0.0001 for significance. (E, F, G) Correlation between *NTRK2* and *EGFR* expression levels in the TCGA GBM dataset. Pearson correlation coefficients and their *p* values were calculated using GraphPad prism. Trend lines were determined by the linear regression model. Overall survival probability in a set of 523 samples from the TCGA GBM cohort. (H) all samples and (I) EGFR amplified samples. Patients were grouped according to low or high expression of *NTRK2*. Survival distribution was estimated according to the Kaplan-Meier method using median cut-off selection and log-rank statistics.

### Correlations between *NTRK2* and *EGFR* expression in GBM tumors

To analyze whether the *NTRK2* expression correlates with *EGFR* levels, we performed correlation analysis across samples registered with TCGA dataset. We detected a weak correlation between *NTRK2* and *EGFR* expression when considering all GBM samples together (Fig. 1E, correlation = 0.1603, *p =* 0.0002) and G-CIMP GBM samples (Fig. 1G, correlation = 0.1869, *p =* 0.0001). A moderate correlation between *NTRK2* and *EGFR* expression was found in G-CIMP+ samples (Fig. 1F, correlation = 0.3443, *p =* 0.0221).

We also determined the relationship between *NTRK2* expression and patient overall survival (Fig. 1H), using median of *NTRK2* expression as a cutoff between low and high expression. We further looked at the effect of *NTRK2* expression in patients stratified by *EGFR* amplification (Fig. 1I). Survival of GBM patients was not significantly different when tumors with high and low *NTRK2* expression levels were compared.

### Selective inhibition of TrkB and EGFR decreases GBM cell viability

To examine the effects of TrkB and EGFR inhibition on cell viability, GBM cells were treated with ANA-12 and AG 1478 alone or in combination and viability was assessed by the trypan blue exclusion method. Time course analysis showed that both single treatments decreased A172 and U87MG cell viability when compared with control cells treated with vehicle (*p <* 0.001). The effect was observed in A172 cells after 48 hours of treatment with ANA-12 at 10 μM or AG1478 at 30 μM (Fig. 2A, 2B). A similar effect was observed in U87MG cells after 24 hours of treatment with the dose of 10 μM of ANA-12 or AG1478 (Fig. 2C, 2D). We also analyzed whether the reduction of cell viability with ANA-12 or AG1478 were dose-dependent. As depicted in Fig. 2, both treatments showed features of dose-dependent effect in A172 and U87MG cells. After time course and dose-response curves analysis, we defined proper doses and time to evaluate the effect of the combination (ANA-12 plus AG1478) on cell viability. When the two drugs were combined the inhibitory effect was more pronounced in A172 cells compared to either IC50-equivalent isolated drugs and in the combination of both drugs in higher doses (see Fig. 2I for detailed results and *p* values for specific comparisons). In contrast, the effect of the combination treatment was not more pronounced than those of each single treatment at equivalent doses in U87MG cells (Fig.2J).

**Fig 2.**
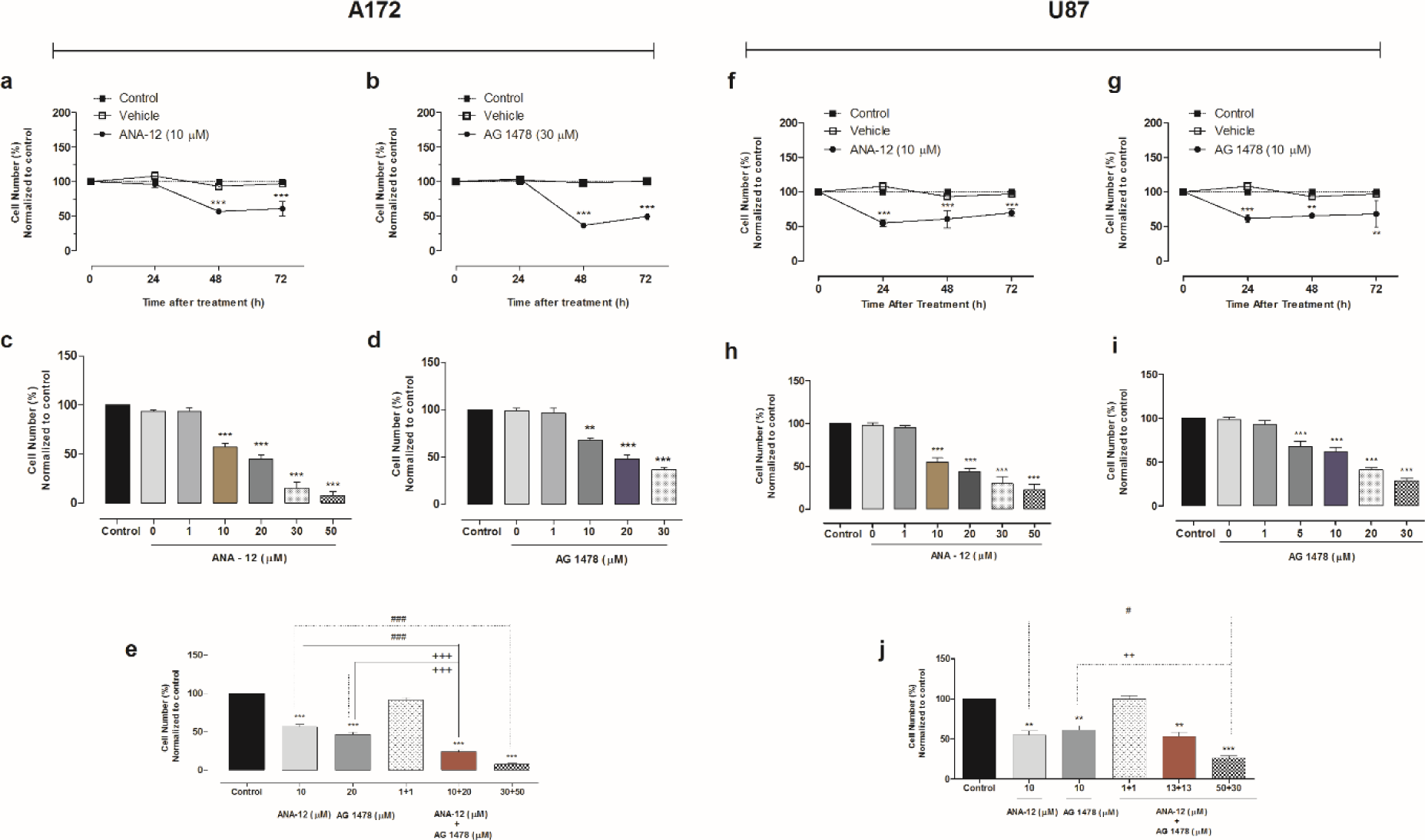
Inhibition of TrkB and EGFR alone or in combination reduces human GBM cell viability. Time course analysis of cell viability, by trypan blue cell counting, were performed after 24, 48 and 72 h of exposure to ANA-12 or AG 1478 exposure in A172 (A, B) and U87MG (C, D) cells. Dose-response curves were evaluated by trypan blue cell counting after treatment with increasing concentrations of ANA-12 (1-50 µM) or AG 1478 (1-30 µM) for 48 h in A172 cells (E, F) and 24 h in U87MG cells (G, H). The drug vehicles (DMSO or EtOH) served as controls. Dose-response curves after combined treatment with ANA-12 and AG 1478 were evaluated after 48 h of drug exposure in A172 cells (I) and 24 h in U87MG cells (J). Data are expressed by mean ± SEM and represent three independent experiments \**p* < 0.05, \*\**p* < 0.01, \*\*\**p* < 0.001, \*\*\*\**p* < 0.0001 compared to control cells; # *p* < 0.05 and ### *p* < 0.001 compared to ANA-12; ++ *p* < 0.01 and +++ *p* < 0.001 compared with AG 1478; one-way ANOVA followed by Tukey tests for multiple comparisons).

### Pharmacological interactions between TrkB and EGFR inhibitors

In order to evaluate drug interaction effects, IC50-values were calculated from the effects seen in the cytotoxicity assay (Fig. 3). ANA-12 had IC50 values of 10.0 (7-14) μM and 13.85 (11-17) μM for A172 and U87MG cells, respectively. IC50 values of AG1478 were 20.0 (16-21) μM for A172 cells and 13.26 (11-16) μM for U87MG cells (Fig. 3A). Pharmacological interactions of the combined treatment with ANA-12 and AG1478 were investigated using cytotoxicity as the chosen outcome and evaluated by Chou-Talalay method (Chou and Talalay 1984). Synergism, addition and antagonism for drug combinations was defined as CI < 0.9, CI = 0.9-1.1 and CI > 1.1, respectively. We observed synergy in A172 cells (CI = 0.75). However, in U87MG cells the inhibitors presented antagonism, as shown by the combination index value of 2.2 (Fig. 3B).

**Fig 3.**
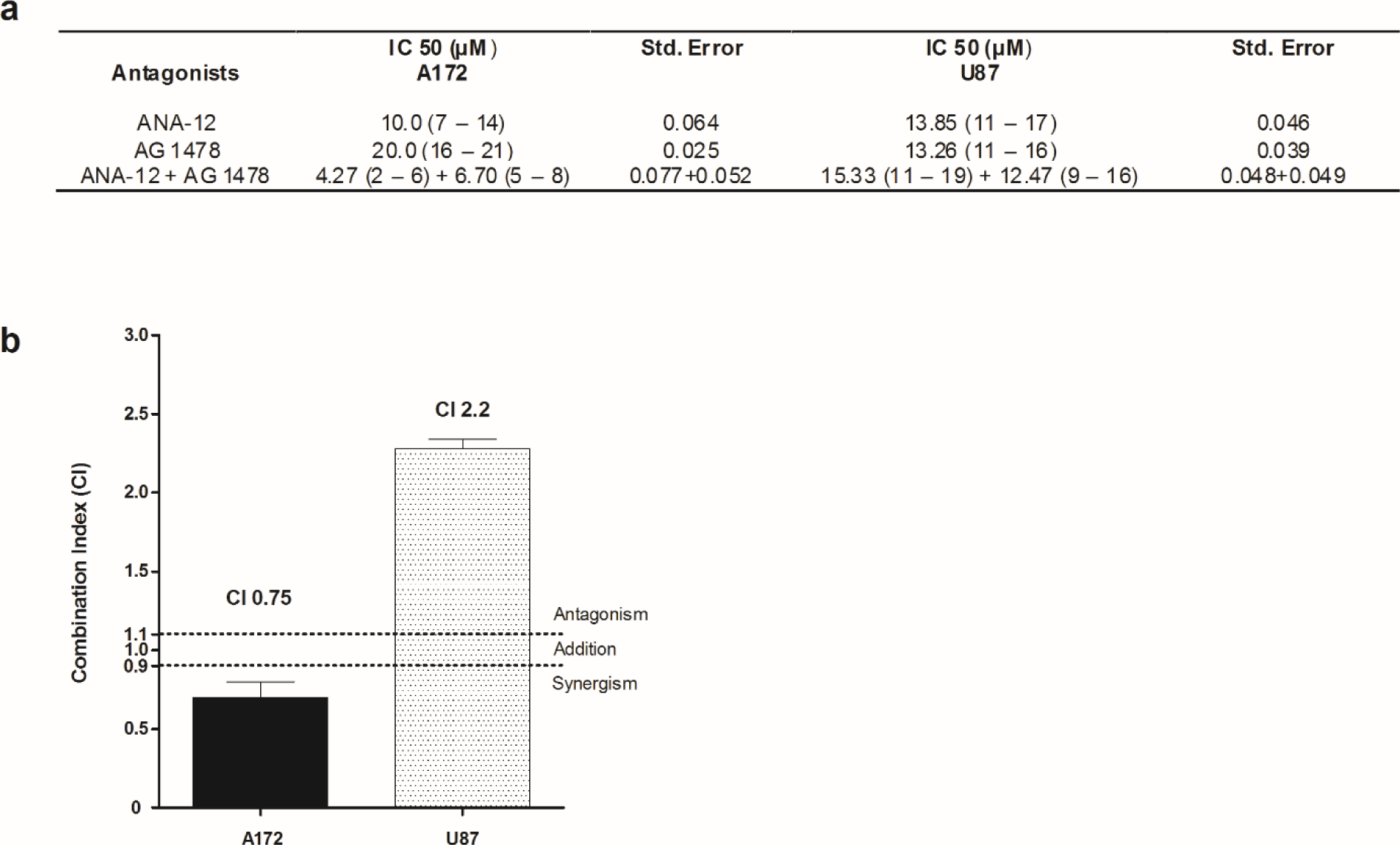
Synergistic effect after combined inhibition of TrkB and EGFR in A172 GBM cells. A172 and U87MG cell lines were treated with varying concentrations of ANA-12 and AG 1478 alone or in combination. The IC50-values were calculated from the dose-response curves after different exposure times (48 h for A172 and 24 h for U87MG cells) and expressed with their respective 95% confidence intervals and summarized in the table (A). The combination index (CI) was determined by the method of Chou-Talalay [20] and data are presented as mean ± SEM (B).

### TrkB and EGFR inhibitors induce changes in cell cycle features of GBM cells

Effects on cell cycle were evaluated by flow cytometry after treatment with ANA-12 and AG 1478 alone or in combination in GBM cells. As shown in Fig. 4, the TrkB inhibitor induced a significant reduction in S phase, which starts at the dose of 10 μM and persists at a dose of 20 μM in A172 cells (Fig. 4A *p <* 0.001 and *p <* 0.0001, respectively). An increase in cells in the S phase after TrkB inhibition was observed in U87MG cells at the dose of 50 μM (*p <* 0.05, Fig. 4D). Treatment with 30 μM of AG 1478 significantly increased the number of A172 and U87MG cells in G0/G1 phase when compared with control cells (Fig. 4B, 4E, *p <* 0.01 and *p <* 0.05, respectively). The combined treatment lead to a combination of effects leading to both accumulation of cells in G0/G1 and reduction of S phase in A172 cells (Fig. 4C *p <* 0.05 and *p* < 0.01). The combined treatment did not cause significant changes in the cell cycle of U87MG cells; however, we could observe a small percentage of cells in Sub-G1 phase when compared with controls (Fig. 4F, *p <* 0.05).

**Fig 4.**
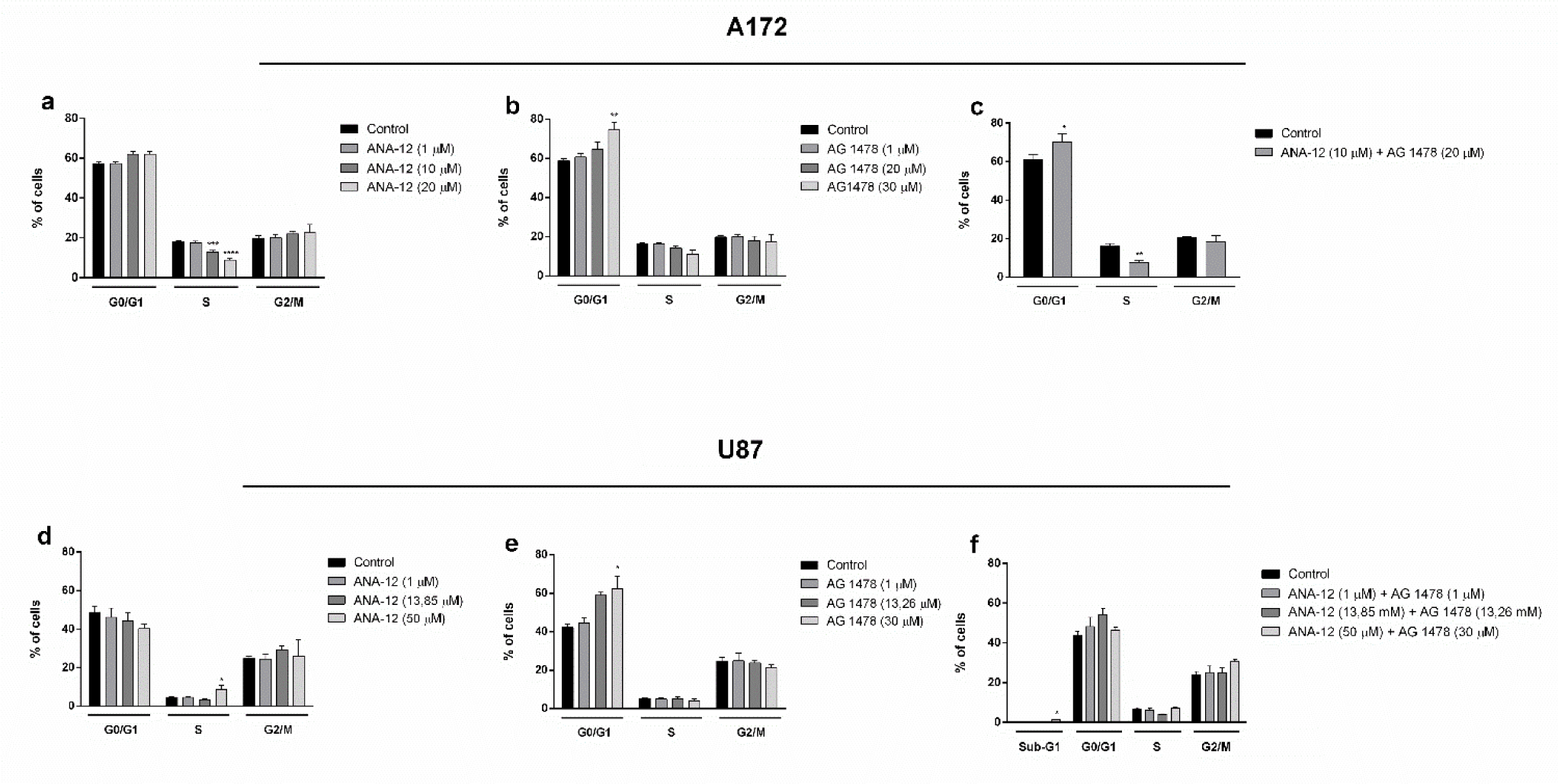
Effects of EGRF and TrkB inhibition on GBM cell cycle. Cells were exposed to ANA-12, AG 1478 or ANA-12 plus AG 1478 for 24 h, and the percentages of cells in G0/G1, S and G2/M phases of the cell cycle were evaluated. Two wells were assigned to each treatment and the experiments were repeated at least three times. Data are presented as mean ± SEM of the percentage of cells in each phase. * *p* < 0.05; ** *p* < 0.01; *** *p* < 0.001 and **** *p* < 0.0001 compared to the respective control group, one-way ANOVA followed by the Dunnett test (**a**,**b**).

### Long-term effects of TrkB and EGFR inhibition in a GBM xenograft mouse model

To explore the roles of EGFR and TrkB during tumorigenesis, we used a preclinical subcutaneous xenograft GBM model. U87MG cells were treated with ANA-12, AG 1478, ANA-12 plus AG 1478 and DMSO or EtOH. After 24 h, the animals were randomized and viable pre-treated U87MG cells were injected subcutaneously into the right flank of mice (Fig. 5A). Fig. 5B illustrates tumor volume size across different days. Tumor size was significantly different in day 15 in all treatment groups when compared with controls given vehicle. Mice that received cells treated with ANA-12, alone or combined with AG 1478, also showed smaller tumors at day 31. The apparent reduction in tumor size in all drug-treated groups compared to controls at day 45 did not reach statistical significance (Fig. 5C). When tumors reached 800-100 mm^3^ the mice were euthanized, and the tumors were excised, and weight and volume were determined. *Ex-vivo* tumor analysis showed an apparent reduction in tumor volume when compared treatment groups with control, however, these differences did not reach statistical significance (Fig. 5D). Moreover, there were no statistically significant differences among groups regarding tumor weight (Fig. 5E). We also evaluated possible differences in survival comparing treatment groups with control group. Although the median survival days of the control group (38 days) was lower when compared to treatment groups (ANA-12, 45 days; AG 1478, 41 days; ANA-12+AG 1478, 44 days), no statistically differences were observed (Fig. 5F).

**Fig 5.**
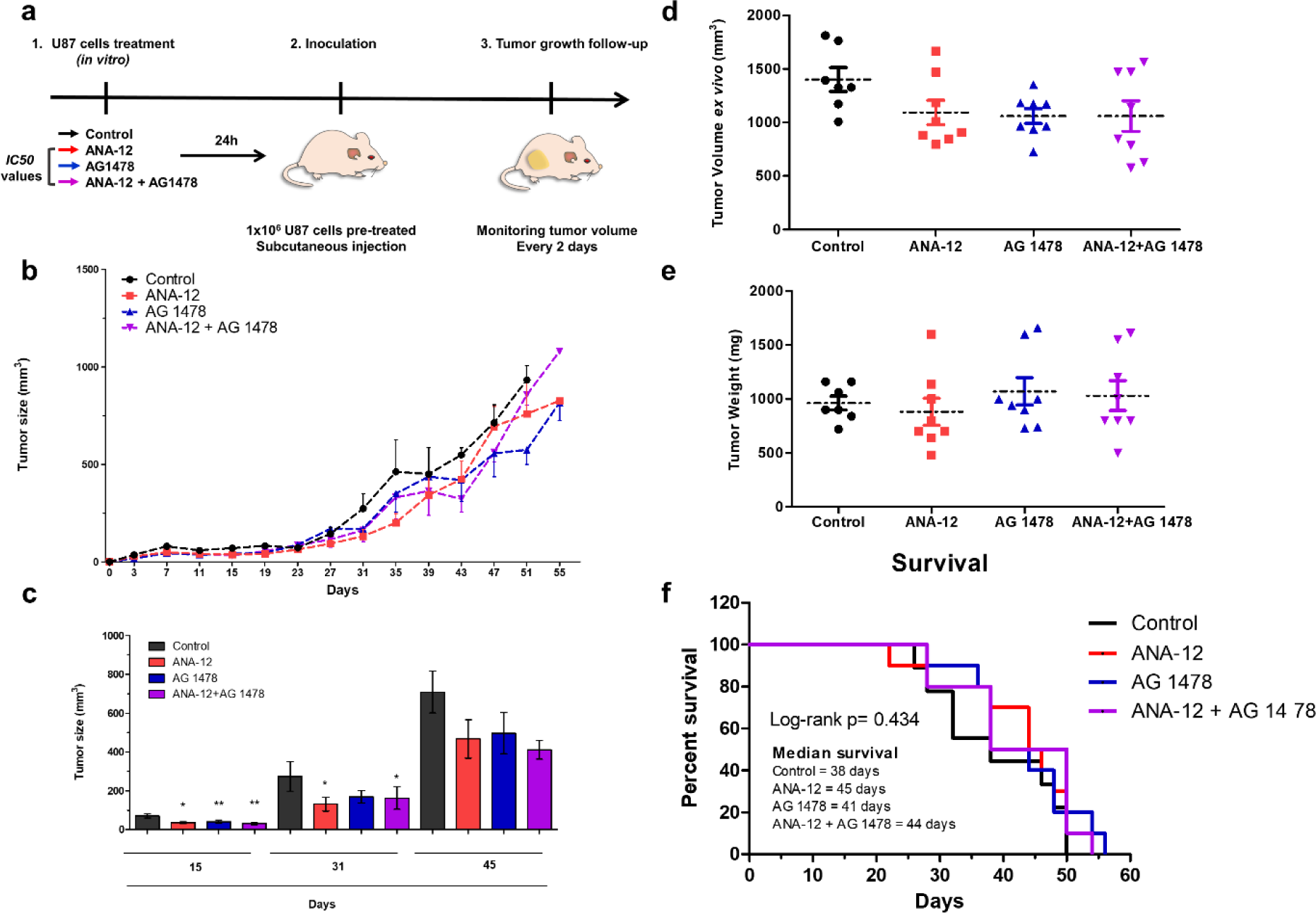
Inhibition of TrkB and EGFR alone or in combination in a subcutaneous GBM xenograft mouse model. U87MG cells were pretreated *in vitro* for 24 h with ANA-12 (13.85 μM), AG-1478 (13.26 μM) or ANA-12 plus AG-1478, and the viable cells were injected into the flanks of nude mice (6-7 mice per group) as shown in the schematic drawing (A). Caliper measurements were used to determine the displayed subcutaneous tumor volume. Mice were euthanized and tumors were excised when volume reached approximately 800-1,000 mm^3^ (B) Tumors were measured every 2 days and volumes were calculated as described in materials and methods section. Tumor growth is represented by tumor volume (mm^3^) at the indicated days; Control (*n=* 7), ANA-12 (*n=* 8), AG 1478 (*n=* 8) and ANA-12 plus AG 1478 (*n=* 8) (C) Tumor growth curve is shown on selected time points of 15, 31 and 45 days to highlight statistical differences (D). Tumor volumes (mm^3^) at the time of tissue harvest. (E) Tumor weight (mg) at the time of tissue harvest are shown in (F) Kaplan-Meier curves presenting percent of mice surviving following tumor implantation. Data are expressed as mean ± SEM (* *p* < 0.05; ** *p* < 0.01). Statistical analysis was performed using one-way ANOVA followed by Tukey’s post-hoc tests.

### Lack of effect of combined TrkB and EGFR inhibition on intracranial GBM tumor growth

We tested the hypothesis that the combined inhibition of TrkB plus EGFR would inhibit the growth of intracranial GBM in mice. Seven days after inoculation of U87MG cells, mice were randomized to receive i.p. injections in a period of 21 days with ANA-12 (1 mg/kg daily plus vehicle every 3 days), AG1478 (10 mg/kg every three days plus vehicle daily), ANA-12 (1 mg/kg daily) plus AG 1478 (10 mg/kg every three days) and vehicle daily. During the treatment period, mice were monitored daily for any signs of sickness, pain or weight loss. The tumor growth was analyzed by ex-vivo brain fluorescent imaging of the different groups using the IVIS system after 29 days after cell inoculation. Tumor fluorescence did not reveal a significant difference among groups (*p =* 0.771; Suppl. Fig. S1).

## Discussion

Gene expression analysis in GBM tumor datasets showed an increased *NTRK2* expression in proneural, G-CIMP^+^ GBMs. Interestingly, significant correlations between *NTRK2* and *EGFR* expression levels were found in GBM, particularly in G-CIMP+ tumors. Previous analyses of TCGA GBMs have shown that G-CIMP+ glioblastomas presented reduced mRNA levels for EGFR due to epigenetic regulation [27]. The proneural G-CIMP phenotype confers a survival advantage for GBM patients [3], and its possible relationship with *NTRK2* expression revealed here warrants further exploration.

One of the main findings in the present study was that the combined inhibition of TrkB and EGFR was more pronounced than either treatment given alone in impairing A172 GBM cell viability. This result is consistent with the view that TrkB inhibition may sensitize cancer cells to the effects of EGFR inhibitors [28]. ANA-12, originally developed as an experimental antidepressant, selectively and efficiently inhibits TrkB by binding to both low-and high-affinity sites on the receptor extracellular domain [21]. AG 1478 is a selective EGFR inhibitor that shares a structural quinazoline main chain with the clinically used EGFR inhibitors gefitinib and erlotinib [20]. The results observed with these two small-molecule compounds in two different mouse GBM xenograft models were less clear. When cells were pretreated with drug treatments before being xenografted into the flanks of nude mice, treatments were able to slow tumor growth only at specific time points (15 and 31 days after cell inoculation), but drug-treated tumors were able to reach sizes comparable to controls by the end of the follow-up period. In addition, treatments did not significantly change tumor progression assessed by fluorescence in mice receiving GBM cells intracranially followed by systemic drug treatments. Given the potency of ANA-12 and AG1478 inhibition of cell viability *in vitro*, we expected to observe pronounced drug effects in *in vivo* models. A possible limitation was measuring fluorescence only at the end of treatment rather than at different time points during treatment. Importantly, EGFR may have a dual role, contributing to tumor progression but also conferring increased DNA damage response and high sensitivity to the inhibitor talazoparib [29]. Thus, differential roles of EGFR signaling could occur under the different experimental conditions we used (for example, regulation by the tumor microenvironment in *in vivo* assays), resulting in contrasting findings. The lack of effects may also be related to the choice of drug doses in our experiments. The drug doses and treatment regimens we used for *in vivo* experiments were based on previous studies that used xenograft models of GBM and other cancer types [22, 24, 25]. It is still possible that, as suggested by our results shown in Fig. 3, the two drugs used show antagonistic activity in U87MG cells. Another point worth mentioning is that, together, the results obtained in our *in vitro* and flank xenograft experiments might suggest that TrkB and EGFR inhibition can delay GBM growth in the short-term, but not after longer delays, when tumors are able to recover full growth.

One issue worthy discussing is the opposite pattern of effects on cytotoxity between the two cell lines. Pharmacological interaction analysis showed synergism in A172 cells but antagonism in U87 MG cells. In terms of genetic profiles, both A172 and U87 MG cells are mutant for both CDKN2A and PTEN. However, U87 MG cells express only wild-type EGFR, whereas A172 cells also express TDM/18-26 mutant EGFR [30, 31]. This biological difference might be crucial to explain the contrasting effects we observed between cell lines. In addition, we have previously observed, in experiments using medulloblastoma cells, that BDNF/TrkB signaling may have opposite effects on cell viability, so TrkB activation can either induce or prevent cytotoxicity depending on drug combinations and experimental conditions [32 - 34]. In fact, BDNF can play opposite actions depending on changes in cellular context. These differential effects may involve distinct isoforms of TrkB expressed in different cell tumor types [34]. This dependency on cellular environment might also be related to the switch from synergism to antagonism we observed when TrkB inhibition was combined with EGFR inhibitor. GBM tumors interact in complex ways with a microenvironment consisting of glial, endothelial, and immune cells. Several signaling molecules, including growth factors, mediate intercellular communication between GBM cells and cells composing the microenvironment [35 - 37]. It is thus possible that the interactions between receptors such as TrkB and EGFR in GBM, as well as the effects of receptor inhibitors, are modulated by biochemical signals from the microenvironment. It is also possible that inhibitors interact directly with receptors on cells in the microenvironment, resulting in changes of drug responses *in vivo* compared to *in vitro* models. These are complex issues that warrant further research in the field of targeted therapies for GBM.

GBM differentiated cells have been shown to secrete BDNF to stimulate TrkB in GBM stem cells and induce them to secrete nerve growth factor inducible (VGF), a neuropeptide that promotes survival and growth of GBM stem-like and differentiated cells [15]. TrkB knockdown or pharmacological inhibition of TrkB hinders BDNF-dependent GBM stem-like cell growth [13]. TrkB-containing exosomes promote the transfer of aggressiveness between GBM cells [38]. EGFR inhibition has been proposed as a therapeutic strategy in GBM and has been tested in preclinical [39 – 41] and clinical [42, 43] studies, with mixed results. One previous study examined the interaction between TrkB and EGFR in experimental GBM and showed that stimulation of neurotrophin signaling can overcome the inhibitory effects of EGFR inhibition on GBM cell growth [13]. The present study is the first one to verify whether the combined inhibition of TrkB and EGFR may effectively reduce GBM growth.

In summary, analysis of TCGA tumors showed that correlations between *NTRK2* and *EGFR* expression can be observed in GBM, and the role of TrkB and EGFR in the proneural, G-CIMP^+^ subtype of GBM should be further investigated by future studies. In addition, our results using cultured cells indicate for the first time the potential of combining TrkB and EGFR inhibition for the treatment of GBM, however we could not observe pronounced effects *in vivo*.

## Acknowledgements

This article has been previously published as a preprint at https://www.biorxiv.org/content/10.1101/2020.02.03.932608v1.

## Research involving human participants and/or animals

Not applicable.

## Informed consent

Not applicable.

