## Supplementary information for "Expression and pharmacological inhibition of TrkB and EGFR in glioblastoma"

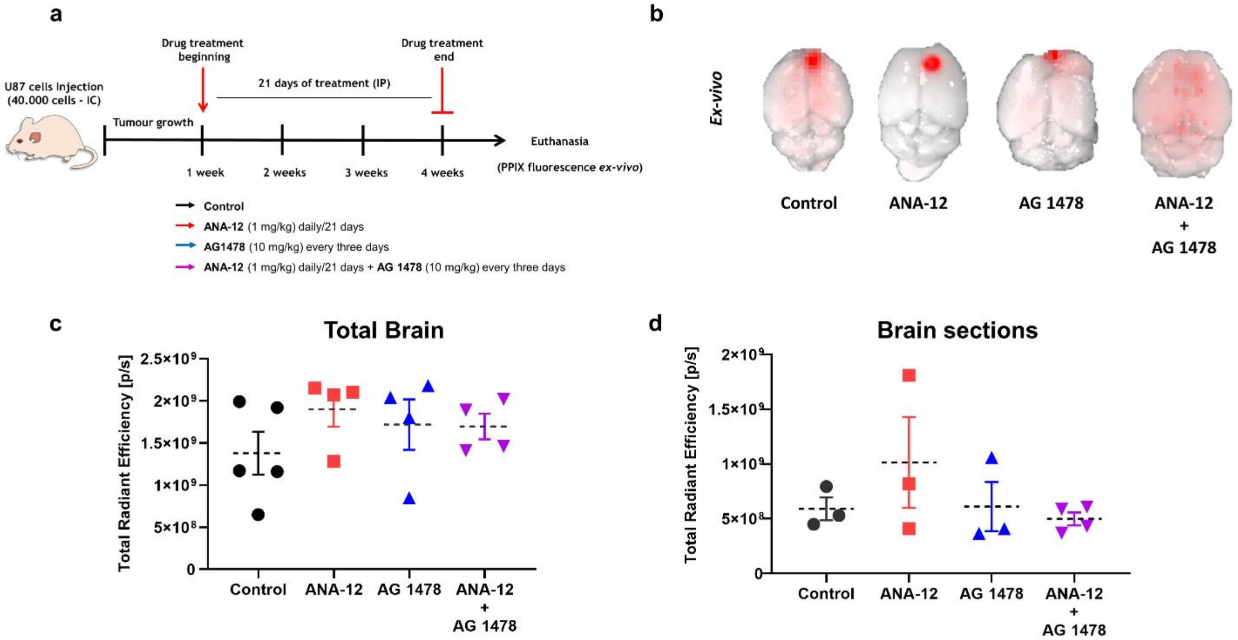


**Supp. Fig. S1** Inhibition of TrkB and EGFR in an intracranial GBM mouse model. (A) A total of 40,000 U87MG cells were intracranially (i.c.) injected into nude mice. Drug treatments started on the seventh day after cell implantation. The animals were randomly divided in 4 groups ($n$=5 per group) to receive intraperitoneal (i.p.) injections for 21 days, and were treated by a blinded investigator with ANA-12 (1 mg/kg daily plus vehicle every 3 days), AG1478 (10 mg/kg every three days plus vehicle daily), ANA-12 (1 mg/kg daily) plus AG 1478 (10 mg/kg/every three days) and vehicle (DMSO) daily. After 21 days of treatment, the animals received an i.p. injection of 5-ALA (50 mg/kg) and after 1 h mice were euthanized by cervical dislocation and the brains were removed to be analyzed  (B) Representative IVIS images of harvested brains acquired at day 29 after transplantation. (C) Corresponding data of 5-ALA (PPIX) radiant efficiency from the whole brain. (D) After performing images of the intact brain, a brain matrix was used to cut sequential 1-mm slices through the region containing the tumor. Slices were imaged using IVIS and fluorescence images were collected. Data are expressed as mean ± SEM. Statistical analyses were performed using one-way ANOVA followed by Tukey's post-hoc tests. No statistical differences were observed between experimental groups.
